# CRISPR-mediated *Stxbp1* gene activation restores Munc18-1 levels in *Stxbp1*-haploinsufficient mice

**DOI:** 10.64898/2026.06.08.730996

**Authors:** Tetsushi Yamagata, Ryu Ikebe, Kenta Kobayashi, Masanari Saito, Yurina Hibi, Kazuhiro Yamakawa, Toshimitsu Suzuki

## Abstract

Mutations in the *syntaxin-binding protein 1* (*STXBP1*) gene, which encodes the presynaptic protein Munc18-1, cause a spectrum of severe epileptic encephalopathies and neurodevelopmental disorders, including Ohtahara syndrome, for which no curative treatment is currently available. Because the disease pathomechanism is thought to be driven by haploinsufficiency, restoring expression of the wild-type allele to physiological levels could provide therapeutic benefit. Here, we evaluated CRISPR-mediated transcriptional activation (CRISPR-ON), based on a dCas9-VPR transcriptional activator, as a strategy to upregulate endogenous *Stxbp1* expression in a *Stxbp1*-haploinsufficient (*Stxbp1*^+/−^) mouse model. Screening of guide RNAs (gRNAs) targeting the *Stxbp1* promoter in Neuro2A cells identified a multiplexed four-gRNA cassette that elevated *Stxbp1* mRNA approximately six-fold. AAV-PHP.eB vectors co-expressing this 4xgRNA cassette and Cre recombinase under the *EF1a* promoter were administered intracerebroventricularly to neonatal *Stxbp1*^+/−^/dCas9-VPR^fl/+^ mice. CRISPR-ON treatment raised brain *Stxbp1* mRNA in *Stxbp1*^+/−^ mice to 1.5-fold the level of wild-type controls, whereas Munc18-1 protein was restored to the wild-type level. In small exploratory cohorts, spike-wave discharge frequency and attack counts tended to be lower in CRISPR-ON–treated than in untreated *Stxbp1*^+/−^ mice; the discharge data, from two animals per group, were not tested statistically, and the difference in attack counts was not significant, so efficacy cannot be judged from the present data. Locomotor activity was normal in untreated mutants and unaffected by treatment; reduced body weight was not rescued. Together, these findings establish that CRISPR-ON can correct *Stxbp1*/Munc18-1 expression in vivo, whereas its effect on disease phenotypes remains to be determined.

## Introduction

Genomic analyses have identified many genes causative for neurodevelopmental disorders and epileptic encephalopathies. Among these, mutations in the *syntaxin-binding protein 1* (*STXBP1*) gene, which encodes Munc18-1, a protein essential for synaptic vesicle fusion and neurotransmitter release [1], are frequently identified in patients with a broad spectrum of epileptic encephalopathies and neurodevelopmental disorders, including the early-onset, severe Ohtahara syndrome [2, 3]. Virtually all patients with *STXBP1* mutations exhibit seizures and intellectual disability, and a subset display autism spectrum features or heightened aggression [4], which complicates care and is associated with later oppositional defiant and conduct disorders [5]. No curative treatment has been established, underscoring the urgent need for therapeutic strategies targeting both seizures and behavioral abnormalities.

The pathomechanism underlying *STXBP1*-related disorders is thought to be primarily driven by haploinsufficiency: Munc18-1 protein levels are reduced to approximately half of wild-type levels in heterozygous *Stxbp1* knockout (*Stxbp1*^+/−^) mice [6], overexpression of mutant alleles does not exacerbate the phenotype [7], and patient-derived mutant Munc18-1 proteins exhibit markedly impaired function and accelerated degradation [2]. Thus, a quantitative reduction in functional Munc18-1 protein appears to be the primary driver of disease onset [8].

Our group has previously demonstrated that *Stxbp1*^+/−^ mice exhibit prominent aggression [6]. Conditional knockout analyses using *Emx1-*Cre (cortical, hippocampal, olfactory bulb, and amygdalar excitatory neurons) [9, 10] and *Vgat-*Cre (all inhibitory neurons) [11] drivers showed that heterozygous loss of *Stxbp1* restricted to either population was insufficient to reproduce this phenotype, suggesting that haploinsufficiency in subcortical excitatory neurons is responsible. We further showed that the aggression was ameliorated by ampakines, which potentiate excitatory neurotransmission, and identified a novel epileptogenic circuit in *STXBP1*- and *SCN2A*-related epilepsies in which reduced cortical excitatory input to striatal fast-spiking inhibitory neurons contributes to seizure onset [6, 12]. Independent groups have reported similar abnormalities in constitutive *Stxbp1*^+/−^ mice and in conditional knockouts using *Vgat-Cre*, *Gad2-Cre*, and *Vglut2-Cre* drivers [7, 13, 14].

We have previously demonstrated the therapeutic potential of CRISPR-mediated activation (CRISPR-ON), in which a nuclease-dead Cas9 (dCas9) fused to the transcriptional activators VP64, p65, and Rta (dCas9-VPR) is guided to a target promoter by a specific guide RNA (gRNA) to enhance endogenous transcription without permanent genomic alterations [15]. In a knock-in mouse model of Dravet syndrome, a pharmacoresistant epileptic encephalopathy caused by haploinsufficiency of *SCN1A* (encoding the voltage-gated sodium channel Nav1.1), CRISPR-ON targeting the *Scn1a* upstream promoter with multiple gRNAs, delivered selectively to inhibitory neurons, restored Nav1.1 protein levels and markedly improved febrile seizures, sudden death, and behavioral abnormalities [16].

Given that *STXBP1*-related disorders share the haploinsufficiency pathomechanism of *SCN1A*-related epilepsies, we hypothesized that CRISPR-ON-mediated upregulation of *Stxbp1* could similarly restore Munc18-1 levels and alleviate disease-associated phenotypes. Here, we designed gRNAs targeting the *Stxbp1* promoter, evaluated their transcriptional activation efficiency both *in vitro* and *in vivo*, and applied this system to *Stxbp1*^+/−^ mice to assess therapeutic efficacy against two key phenotypes: epileptiform spike-wave discharges (SWDs) and excessive aggression in male mice. Our findings show that CRISPR-ON elevates *Stxbp1* mRNA and restores Munc18-1 protein levels *in vivo*, but its phenotypic effect remains inconclusive.

## Results

### Upregulation of *Stxbp1* gene transcription by CRISPR-ON *in vitro*

To establish a CRISPR-ON system targeting *Stxbp1*, we first screened guide RNAs (gRNAs) directed against the murine *Stxbp1* promoter region. Fourteen gRNAs were designed to tile across the promoter (**Figure 1A**), and each gRNA was individually co-transfected with Sp-dCas9-VPR into Neuro2A cells, a mouse neuroblastoma–derived cell line. *Stxbp1* mRNA levels were subsequently quantified by real-time quantitative PCR (qPCR). Each gRNA was compared with both control conditions, untransfected Neuro2A cells and cells transfected with the empty vector (negative control, NC), by one-way ANOVA followed by Dunnett’s post-hoc test performed separately with each control as the reference; all comparisons are reported in **Supplementary Table S4**.

**Figure 1.**
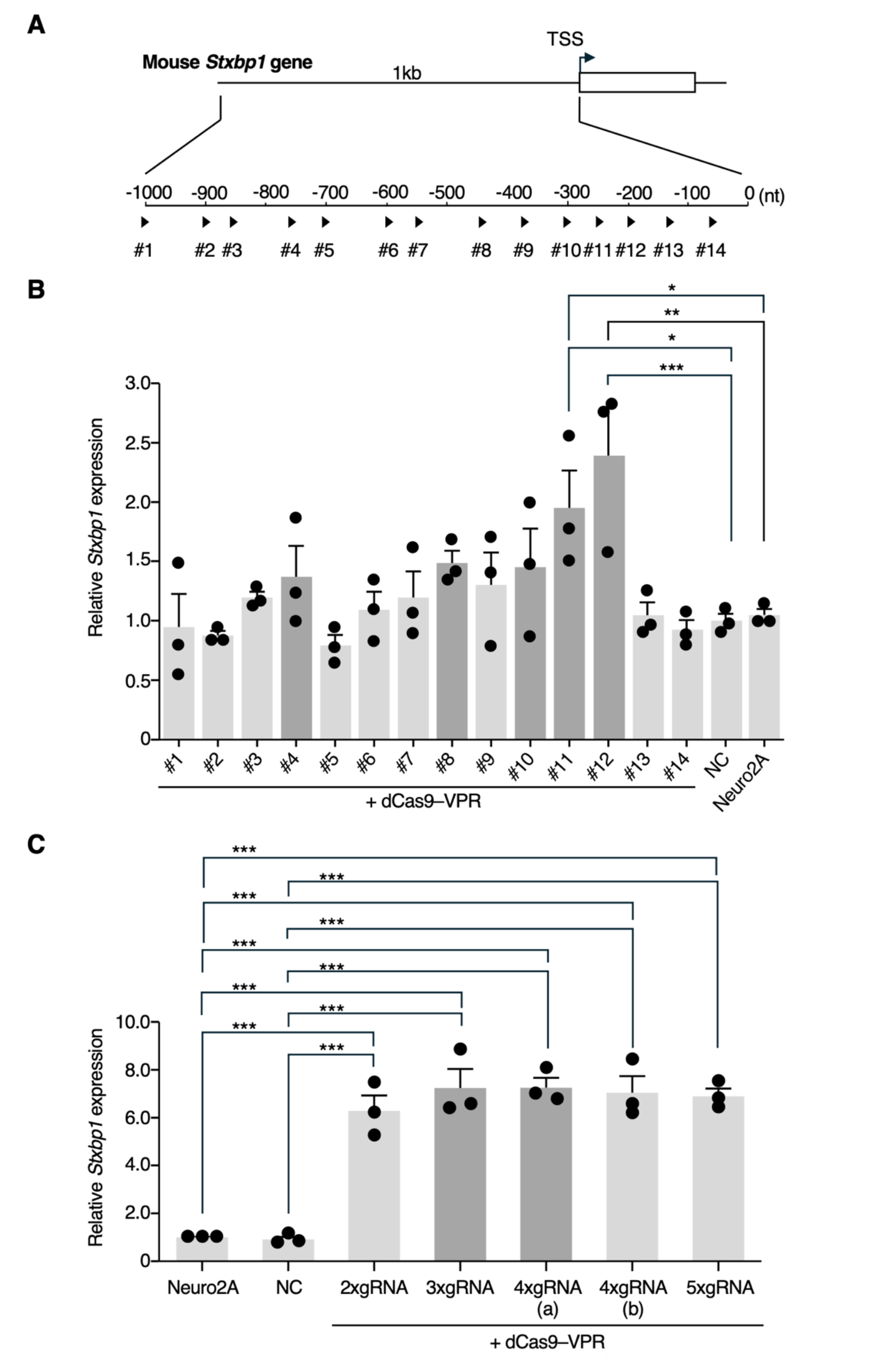
Multiple gRNAs synergistically activate transcription of the endogenous mouse *Stxbp1* gene *in vitro*. **(A)** Schematic of the upstream *Stxbp1* promoter showing the positions of the 14 candidate gRNA sequences selected for screening (#1–#14; arrowheads). The transcription start site (TSS) is indicated by an arrow. **(B)** Real-time qPCR analysis of *Stxbp1* mRNA levels in Neuro2A cells co-transfected with individual gRNAs and dCas9-VPR. Co-transfection of gRNA #11 or #12 with dCas9-VPR significantly increased *Stxbp1* mRNA levels by 1.9- and 2.4-fold, respectively, relative to both control conditions; no other gRNA differed significantly from either control. *Stxbp1* mRNA levels were normalized to those of untransfected Neuro2A cells, and cells transfected with empty vector (MLM3636 plasmid lacking a gRNA target sequence; negative control, NC) served as an additional control. **(C)** Real-time qPCR analysis of *Stxbp1* mRNA levels in Neuro2A cells co-transfected with multiplexed gRNA combinations and dCas9-VPR. Combining multiple gRNAs (two to five per pool) increased *Stxbp1* mRNA expression 6.3- to 7.3-fold. The gRNA pools were as follows: 2xgRNA mix (#11, #12); 3xgRNA mix (#10, #11, #12); 4xgRNA (a) mix (#4, #10, #11, #12); 4xgRNA (b) mix (#8, #10, #11, #12); and 5xgRNA mix (#4, #8, #10, #11, #12). Normalization and the negative control (NC) were as described in (B). In both (B) and (C), n = 3 independent transfections. Statistical comparisons were performed using one-way ANOVA followed by Dunnett’s post-hoc test, carried out separately with untransfected Neuro2A cells and with the NC as the reference control; all comparisons, including those that did not reach significance, are reported in **Supplementary Table S4**, and the tests of model assumptions in **Supplementary Table S6** (\**p* < 0.05, \*\**p* < 0.01, \*\*\**p* < 0.001).

Only gRNAs #11 and #12 differed significantly from either control, yielding 1.9- and 2.4-fold increases in *Stxbp1* mRNA expression, respectively (**Figure 1B**). The remaining twelve gRNAs were not distinguishable from either control, and the two control conditions did not differ from each other, indicating that transfection itself did not alter *Stxbp1* expression. Because previous work has demonstrated that combining multiple gRNAs can synergistically enhance CRISPR-ON activity [16], we next tested whether multiplexing the top-performing gRNAs would further increase *Stxbp1* induction. The five gRNAs with the highest mean activation when tested individually (#4, #8, #10, #11, and #12) were combined into pools of two, three, four, or five gRNAs and co-transfected with Sp-dCas9-VPR into Neuro2A cells. This selection was based on rank order of mean values and was exploratory, since apart from #11 and #12 these gRNAs were not individually distinguishable from the controls. All multiplexed combinations elevated *Stxbp1* mRNA levels by approximately 6- to 7-fold compared with controls, representing a substantial improvement over gRNAs tested individually (**Figure 1C**). With the five gRNAs tested here, activation did not increase in a graded manner with guide number: pools of two to five gRNAs all produced 6.3- to 7.3-fold activation, and no significant differences were detected among them. The choice among them was therefore based on mean activation rather than on a statistical difference: we carried forward the combination with the highest mean, 4xgRNA (a) (#4, #10, #11, #12), and the next highest, 3xgRNA (#10, #11, #12), hereafter referred to as 4xgRNA and 3xgRNA, respectively. To identify an optimal configuration for *in vivo* delivery, we generated AAV constructs combining the 4xgRNA or 3xgRNA cassettes with either an *EF1a*-Cre or *CMV*-Cre, transfected these into Neuro2A cells, and quantified *Stxbp1* mRNA expression. The 4xgRNA construct combined with *EF1a*-Cre gave the highest mean induction, a 6.0-fold increase in *Stxbp1* mRNA levels, although the differences among the construct-containing groups were not statistically significant (**Supplementary Figure S1**). On this basis the 4xgRNA/*EF1a*-Cre configuration was selected for *in vivo* application.

### Generation of double-mutant mice and AAV vectors to establish CRISPR-ON *in vivo*

To verify that the *EF1a* promoter, which produced the highest activation efficiency *in vitro*, would drive sufficient Cre expression in the brain, we administered AAV-PHP.eB-*EF1a*-Cre to ROSA-tdTomato reporter mice via retro-orbital injection and assessed tdTomato fluorescence at 15 weeks of age. Cre-mediated tdTomato expression was observed predominantly in rostral brain regions, including the cerebral cortex, hippocampus, striatum, and thalamus (**Figure 2A**). Given this rostral bias in *EF1a*-driven Cre expression, we sought to extend CRISPR-ON activity to more caudal brain regions by combining *EF1a*-Cre with a *Vglut2*-Cre driver (**Figure 2A**). AAV-PHP.eB particles encoding 4xgRNA together with *EF1a*-Cre, as well as a separate AAV vector encoding *Vglut2*-Cre alone, were generated and co-administered by intracerebroventricular (ICV) injection on postnatal day 1 to *Stxbp1*^+/−^/dCas9-VPR^fl/+^ mice (**Figure 2B**). During the 35-day period following AAV administration, four of five *Stxbp1*^+/−^/dCas9-VPR^fl/+^ mice that received 4xgRNA together with both *EF1a*-Cre and *Vglut2*-Cre died between postnatal weeks 3 and 4, whereas one of five mice that received 4xgRNA with *EF1a*-Cre alone died, as did one of four untreated *Stxbp1*^+/−^ mice (**Figure 2C**). Because natural breeding yielded small and inconsistent litters, each group comprised four to six animals, and no comparison reached statistical significance (*EF1a*-Cre + *Vglut2*-Cre versus saline, *P* = 0.21, Fisher’s exact test). Based on these findings, subsequent experiments employed *EF1a*-Cre alone to drive CRISPR-ON activity, and mice were generated by *in vitro* fertilization and embryo transfer to ensure consistent and adequate cohort sizes. Under this revised protocol, survival of CRISPR-ON–treated *Stxbp1*^+/−^ mice was significantly reduced relative to saline-treated *Stxbp1*^+/−^ mice (7 of 13 versus 0 of 8 died by P35, *P* = 0.018, Fisher’s exact test) and relative to the three control groups combined (*P* < 0.001), although the precise cause remains to be determined (**Figure 2D**).

**Figure 2.**
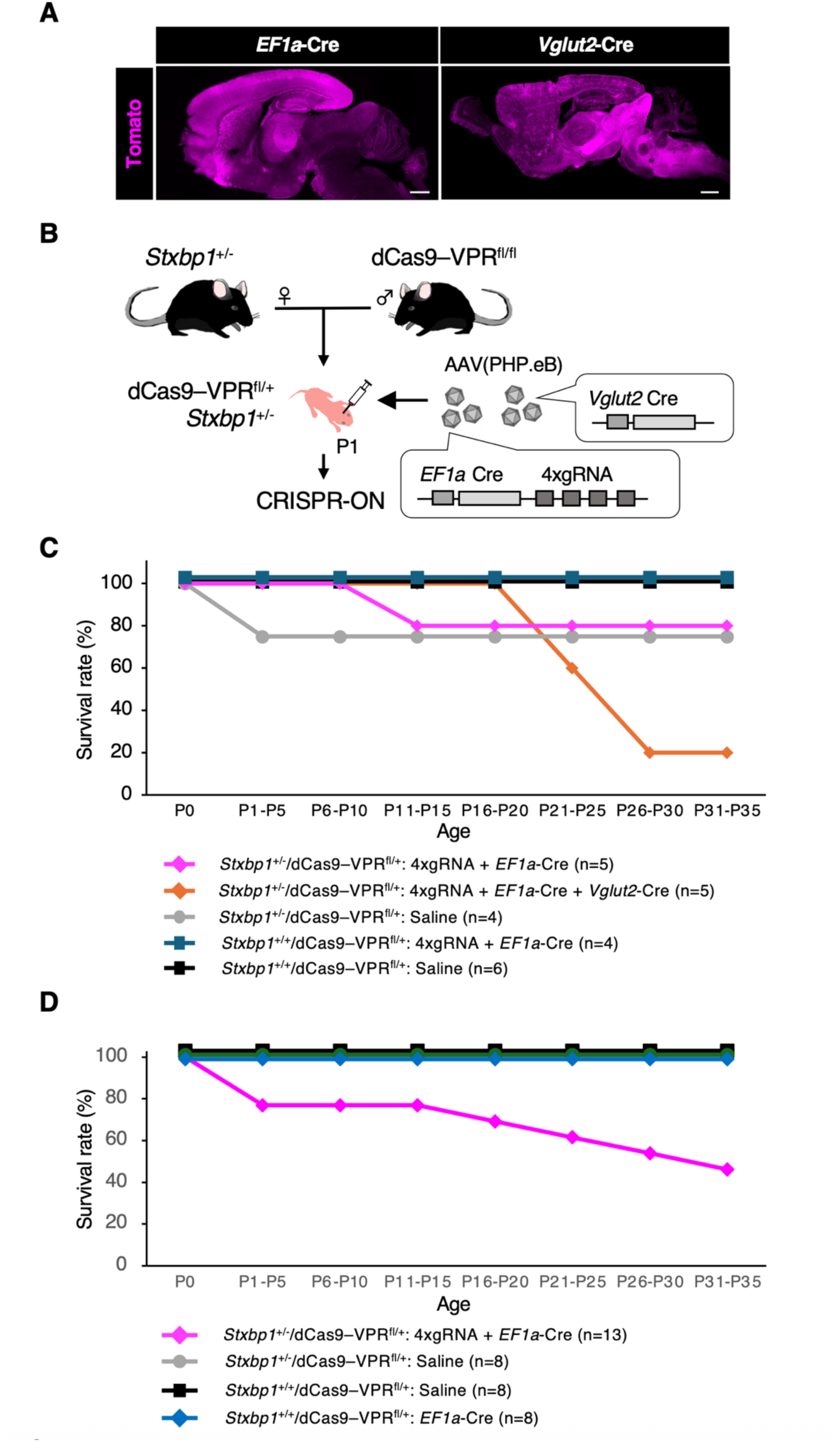
In vivo establishment of CRISPR-ON and its effect on survival in *Stxbp1*-haploinsufficient mice. (**A**) Brain-region-specific Cre expression driven by the *EF1a* or *Vglut2* promoter, visualized by Cre-dependent tdTomato fluorescence (magenta) in ROSA-tdTomato reporter mice. ROSA-tdTomato mice (10 weeks old) received a retro-orbital injection of AAV-PHP.eB-*EF1a* core-Cre (5.6 × 10¹⁰ vg/mouse) or AAV-PHP.eB-*Vglut2*-mCherry-P2A-Cre (4.5 × 10¹⁰ vg/mouse), and brains were fixed 5 weeks later (15 weeks of age). Under the *EF1a* promoter, Cre expression was predominantly rostral, including the cerebral cortex, hippocampus, striatum, and thalamus (left). Under the *Vglut2* promoter, Cre expression was predominantly ventral and caudal, including the thalamus, hypothalamus, pons, and medulla (right). (**B**) Experimental design for establishing CRISPR-ON *in vivo*. *Stxbp1*^+/−^ mice were crossed with dCas9-VPR^fl/fl^ mice, and the resulting F1 offspring received AAV-PHP.eB vectors delivering Cre recombinase and gRNAs to drive CRISPR-ON. (**C**) Combined *EF1a*-Cre and *Vglut2*-Cre CRISPR-ON reduced survival. Survival over the 35-day rearing period following intracerebroventricular AAV administration on postnatal day (P) 1 in offspring. Survival declined between P16 and P30 in the group receiving combined *EF1a*-Cre and *Vglut2*-Cre CRISPR-ON (*Stxbp1*^+/−^/dCas9-VPR^fl/+^ treated with *EF1a*-Cre + 4xgRNA + *Vglut2*-Cre). *n* = number of animals used (indicated below the graph). (**D**) CRISPR-ON treatment reduced survival. Survival to postnatal day (P) 35 in CRISPR-ON–treated mice. Survival declined during P1–P5 and after P15 only in the CRISPR-ON–treated group (*Stxbp1*^+/−^/dCas9-VPR^fl/+^ treated with *EF1a*-Cre + 4xgRNA). n = number of animals used (indicated below the graph).

### CRISPR-ON elevates *Stxbp1* mRNA and restores Munc18-1 protein in the *Stxbp1*-haploinsufficient mouse brain

To determine whether CRISPR-ON could elevate *Stxbp1* expression *in vivo*, brain *Stxbp1* mRNA and Munc18-1 protein levels were assessed at 5 weeks of age in CRISPR-ON–treated *Stxbp1*^+/−^ mice. Real-time qPCR analysis showed that CRISPR-ON treatment raised *Stxbp1* mRNA in *Stxbp1*^+/−^ mice from approximately half the wild-type level to 1.5-fold that level, significantly exceeding untreated wild-type controls (*P* = 0.002; **Figure 3A**). By contrast, Western blot analysis showed that Munc18-1 protein in CRISPR-ON–treated *Stxbp1*^+/−^ mice was restored from approximately half the wild-type level and did not differ significantly from wild-type controls (**Figure 3B** and **Supplementary Figure S2**). The same dissociation was apparent in treated wild-type mice, in which a 2.8-fold increase in *Stxbp1* mRNA was accompanied by only a 1.4-fold increase in Munc18-1 protein.

**Figure 3.**
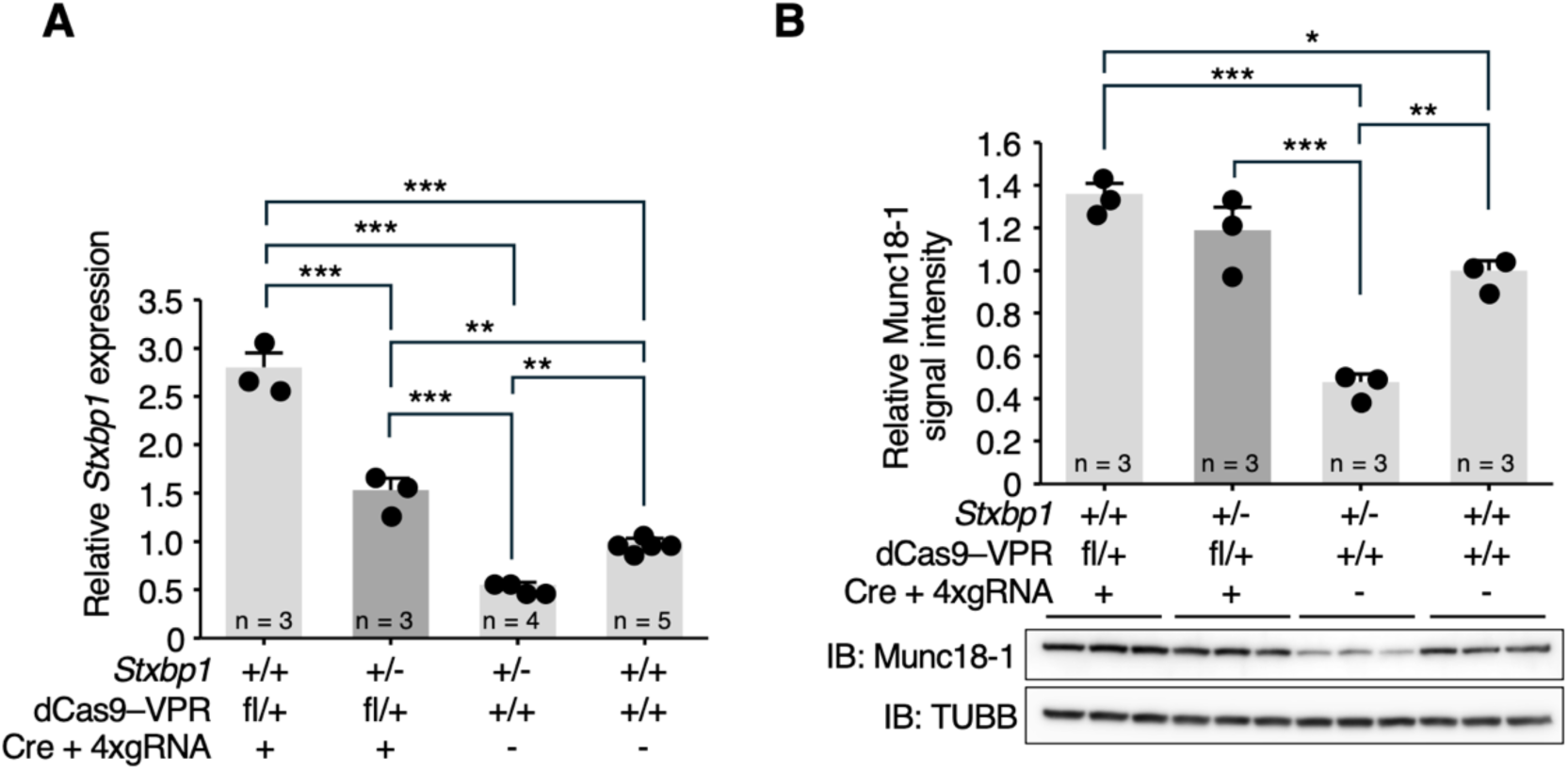
Upregulation of *Stxbp1* mRNA and Munc18-1 protein in the brain of CRISPR-ON–treated *Stxbp1*+/− mice. **(A)** Real-time qPCR quantification of *Stxbp1* mRNA levels in brain hemispheres of CRISPR-ON–treated mice (5 weeks of age). In untreated *Stxbp1*^+/−^ mice, *Stxbp1* mRNA levels were reduced to approximately half of those in control mice (*Stxbp1*^+/+^/dCas9-VPR^+/+^), whereas CRISPR-ON treatment raised them to 1.5-fold the control level. In CRISPR-ON–treated wild-type mice (*Stxbp1*^+/+^/dCas9-VPR^fl/+^/Cre + 4xgRNA), *Stxbp1* mRNA levels were elevated to 2.8-fold those of untreated control mice. The “Cre + 4xgRNA” indicates administration of AAV-PHP.eB-*EF1a*-Cre-4xgRNA, and “−” indicates administration of saline. **(B)** Western blot analysis of Munc18-1 protein levels in brain hemispheres of CRISPR-ON– treated mice (5 weeks of age). In untreated *Stxbp1*^+/−^ mice, Munc18-1 levels were reduced to approximately half of those in control mice (*Stxbp1*^+/+^/dCas9-VPR^+/+^), whereas CRISPR-ON treatment restored expression to control levels. Representative immunoblots for Munc18-1 and the loading control β-tubulin (TUBB) are shown at the bottom. In both (A) and (B), the number of animals used (n) is indicated within the graph. Statistical comparisons were performed using one-way ANOVA followed by Tukey’s post-hoc test; all comparisons are reported in **Supplementary Table S5**, and the tests of model assumptions in **Supplementary Table S6** (\**p* < 0.05, \*\**p* < 0.01, \*\*\**p* < 0.001).

### Exploratory assessment of spike-wave discharges, aggression, and locomotor activity in CRISPR-ON–treated *Stxbp1*^+/−^ mice

We next examined whether this correction of Munc18-1 levels translated into amelioration of the epileptic and behavioral phenotypes characteristic of *Stxbp1* haploinsufficiency. To assess the effect of CRISPR-ON on epileptiform activity, we performed electrocorticography (ECoG) recordings and quantified the occurrence of spike-wave discharges (SWDs), a hallmark electrographic abnormality observed in *Stxbp1*^+/−^ mice (representative traces shown in **Figure 4A**). Because only two animals per group could be recorded, no statistical comparison was performed. In these animals, the mean SWD frequency was lower in CRISPR-ON–treated than in untreated *Stxbp1*^+/−^ mice (**Figure 4B**); however, the highest value in the treated group was close to the lowest value in the untreated group, and with two animals per group this difference cannot be distinguished from inter-animal variability. This observation is therefore descriptive only. While CRISPR-ON treatment may improve this electrographic phenotype, its efficacy cannot be judged on the basis of the present data.

**Figure 4.**
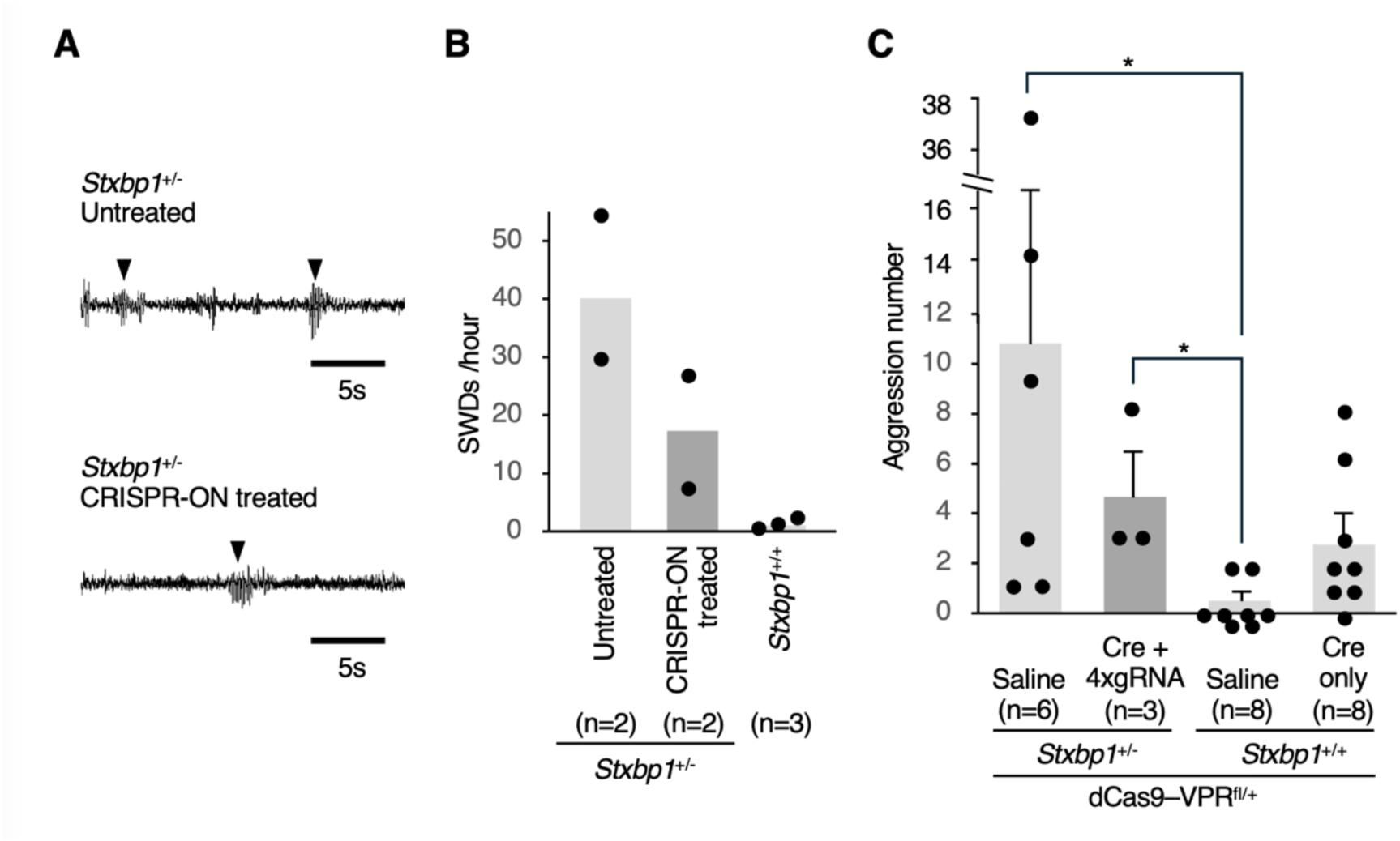
Exploratory assessment of epileptiform discharges and aggression in CRISPR-ON–treated *Stxbp1*^+/−^ mice. **(A)** Representative electrocorticographic (ECoG) traces of spike-wave discharges (SWDs) in CRISPR-ON–untreated and – treated *Stxbp1*^+/−^ mice. **(B)** Quantification of SWD frequency (10–13 weeks of age), counted over a 24-h period (08:00 to 08:00 the following day). Mean SWD frequency was lower in CRISPR-ON–treated mice (*Stxbp1*^+/−^/dCas9-VPR^fl/+^ administered AAV-*EF1a*-Cre + 4xgRNA) than in untreated mice (*Stxbp1*^+/−^/dCas9-VPR^fl/+^ administered saline). Control mice (*Stxbp1*^+/+^) showed almost no SWDs. Because only two animals per group were recorded, no statistical analysis was performed and these data are descriptive only. **(C)** Aggressive behavior assessed by the resident-intruder test (16 weeks of age; 10-min sessions). Attack frequency varied widely among untreated *Stxbp1*^+/−^ mice, and the values of the three CRISPR-ON–treated *Stxbp1*^+/−^ mice all lay within the range of the untreated group; no significant difference was detected between these two groups. Both *Stxbp1*^+/−^ groups showed significantly more attacks than control mice (*Stxbp1*^+/+^/dCas9-VPR^fl/+^ administered saline). Mice expressing dCas9-VPR alone (Cre only) showed numerically higher attack frequencies than control mice, but this difference was not statistically significant. Statistical comparisons were performed using the Kruskal-Wallis test (\**p* < 0.05). Given the small group sizes, these behavioral comparisons should be regarded as exploratory. (B, C) The number of animals used (n) is indicated below the graph.

We then evaluated aggressive behavior using the resident-intruder test. The number of attack episodes was highly variable among untreated *Stxbp1*^+/−^ mice (range, 1-37 attacks per 10-min session; n = 6), and the values of the three CRISPR-ON–treated *Stxbp1*^+/−^ mice available for testing all fell within this range (Figure 4C). No significant difference was detected between untreated and CRISPR-ON–treated *Stxbp1*^+/−^ mice, whereas both groups showed significantly more attacks than *Stxbp1*^+/+^ controls (Kruskal-Wallis test with post hoc comparisons). These data therefore do not provide evidence that CRISPR-ON treatment modified aggressive behavior, and this comparison should be regarded as exploratory and underpowered. Wild-type mice that received *EF1a*-Cre and therefore expressed dCas9-VPR alone in the brain showed numerically higher attack frequencies than untreated wild-type mice (**Figure 4C**), raising the possibility that dCas9-VPR expression itself may affect aggressive behavior; this difference, however, also did not reach statistical significance, and the underlying mechanism remains to be determined. No significant differences were detected among groups in the frequency of non-aggressive behaviors (**Supplementary Figure S3**).

To further characterize behavioral outcomes, we assessed locomotor activity and anxiety-like behavior in a novel environment using the open field test. Untreated *Stxbp1*^+/−^ mice did not differ significantly from wild-type controls in total distance traveled, time spent in the center zone, or average locomotor velocity, indicating that locomotor activity is not impaired in this model; CRISPR-ON–treated *Stxbp1*^+/−^ mice likewise did not differ from untreated *Stxbp1*^+/−^ mice in any of these measures (**Supplementary Figure S4A–C**). However, wild-type mice expressing dCas9-VPR alone in the brain following *EF1a*-Cre administration exhibited significant increases in distance traveled, center-zone occupancy, and locomotor velocity compared with untreated wild-type controls (**Supplementary Figure S4A–C**). These findings suggest that expression of dCas9-VPR alone may induce hyperactivity, although the precise underlying mechanism remains unclear.

Finally, because reduced body weight has previously been reported as a phenotype of *Stxbp1*^+/−^ mice [6, 14], we measured body weight across experimental groups. Both CRISPR-ON–treated and untreated *Stxbp1*^+/−^ mice showed reduced body weight compared with wild-type controls, and CRISPR-ON treatment did not rescue this phenotype (**Supplementary Figure S4D**).

## Discussion

In the present study, we show that CRISPR-mediated transcriptional activation (CRISPR-ON) of the endogenous *Stxbp1* gene restores Munc18-1 protein levels in the brain of *Stxbp1*-haploinsufficient mice. Although *Stxbp1* mRNA in treated *Stxbp1*^+/−^ mice exceeded the wild-type level by 1.5-fold, Munc18-1 protein did not differ from wild type, and in treated wild-type mice a 2.8-fold increase in mRNA produced only a 1.4-fold increase in protein. Munc18-1 abundance therefore appears to be buffered post-transcriptionally. This may limit the risk of protein overexpression when transcription is driven above the physiological range, but it also means that transcript levels overestimate the degree of protein correction, and that the magnitude of transcriptional activation is not a reliable proxy for the magnitude of protein change. In a small exploratory cohort, treated mice showed a lower frequency of SWDs and fewer attacks than untreated mice; the electrocorticographic groups comprised two animals each and were not tested statistically, and the difference in attack counts was not significant. We therefore cannot conclude that CRISPR-ON ameliorates these phenotypes, and the following discussion of the possible cellular basis of these observations should be read as hypothesis-generating rather than as an account of a demonstrated therapeutic effect.

SWD frequency tended to be lower in CRISPR-ON–treated *Stxbp1*^+/−^ mice, although, as noted above, this observation is based on two animals per group and is not statistically supported. Should this trend be confirmed in adequately powered cohorts, our previous work using cell-type-specific conditional knockout (cKO) mice would suggest a plausible cellular basis. Mice with *Stxbp1* deletion in excitatory neurons via *Emx1-Cre*, or in cortical layer 5 excitatory neurons (presumably corticostriatal projection neurons) via *Trpc4-Cre*, exhibit prominent SWDs, whereas mice with inhibitory neuron–restricted deletion via *Vgat-Cre* do not [12]. Because the *EF1a* promoter used here drives Cre expression broadly across neuronal populations,

CRISPR-ON-mediated upregulation of *Stxbp1* likely occurred in *Emx1*- and *Trpc4*-positive excitatory neurons, among others. The most plausible basis would then be restoration of *Stxbp1* expression in excitatory neurons of rostral cortical regions rather than in inhibitory neurons. Any incomplete effect would be consistent with suboptimal transduction efficiency of the AAV-PHP.eB vectors used to deliver Cre and gRNAs *in vivo*. Notably, despite the rostral bias of *EF1a*-driven Cre activity, whole-brain *Stxbp1* mRNA exceeded and Munc18-1 protein reached wild-type levels. This apparent discrepancy suggests that CRISPR-ON achieves robust per-cell transcriptional activation in transduced neurons, even when the proportion of transduced cells across the brain is incomplete. Accordingly, enhancing AAV transduction efficiency in neurons that endogenously express *Stxbp1* will be an important step in testing whether a more complete effect on SWDs can be achieved.

Aggressive behavior in the resident-intruder test was highly variable among untreated *Stxbp1*^+/−^ mice, and the small number of treated animals precludes any conclusion as to whether this phenotype was modified. The cellular basis of this phenotype appears more complex than that of SWDs: in our previous studies, *Stxbp1* haploinsufficiency restricted to excitatory neurons via *Emx1-Cre*, or to inhibitory neurons via *Vgat-Cre*, each failed to recapitulate the aggressive phenotype of constitutive *Stxbp1*^+/−^ mice [6], suggesting that haploinsufficiency in a single neuronal population is insufficient to elicit aggression and that concurrent dysfunction of excitatory and inhibitory circuits is required. Whether a strategy such as CRISPR-ON, which simultaneously elevates *Stxbp1* expression in both populations within transduced regions, can ameliorate aggression, and whether rostral circuits potentially involving the cortex, hippocampus, striatum, and/or amygdala contribute to this phenotype, will need to be addressed in adequately powered cohorts.

CRISPR-ON treatment did not ameliorate the reduced body weight that has been consistently reported in *Stxbp1*^+/−^ mice [6, 13]. Because *EF1a*-Cre activity in our hands was largely restricted to rostral regions, this negative result suggests that the circuits regulating body weight in *Stxbp1*^+/−^ mice reside in more caudal brain structures, including the hypothalamus, midbrain, pons, and medulla. Extending CRISPR-ON activity into these caudal regions would be required to test this possibility.

Sudden unexpected death is well documented in patients with *STXBP1* mutations [17], and reduced survival has likewise been reported in *Stxbp1*^+/−^ mice [7, 13]. In our hands, the survival of *Stxbp1*^+/−^ mice treated with *EF1a*-Cre-driven CRISPR-ON was comparable to that of untreated *Stxbp1*^+/−^ mice when animals were obtained by natural breeding, but was lower than that of untreated *Stxbp1*^+/−^ controls when cohorts were generated by *in vitro* fertilization (IVF) and embryo transfer. The apparent discrepancy between the two cohorts is most simply explained by the size of the natural-breeding experiment. Mortality in CRISPR-ON–treated *Stxbp1*^+/−^ mice was one of five animals in that cohort and seven of thirteen in the in vitro fertilization cohort, a difference that is not statistically significant, and the confidence interval around the former estimate encompasses the latter. Background mortality also differed, with one of four saline-treated *Stxbp1*^+/−^ mice dying in the natural-breeding cohort and no control animal dying in the in vitro fertilization cohort, indicating generally poorer survival in the smaller cohort. Other contributions cannot be excluded. The two cohorts were treated with different preparations of the AAV vector, and batch-to-batch variation in titre or in the proportion of empty capsids could alter the effective dose despite the same nominal dose being administered. In vitro fertilization and embryo transfer are unlikely to be responsible, since the same procedure was applied to the control groups of that cohort, which showed no excess mortality. Drift in genetic background cannot be formally excluded, although both cohorts derived from the same colony. The reduced survival observed in the IVF-derived cohort can be interpreted in light of the regional bias of *EF1a*-Cre activity. Two observations are key: (i) *EF1a*-Cre drove CRISPR-ON activity preferentially in rostral regions; and (ii) elevation of rostral *Stxbp1* in wild-type mice via the same construct did not reduce survival. Together, these observations raise the possibility that the survival cost in CRISPR-ON–treated *Stxbp1*^+/−^ mice does not arise from rostral overexpression *per se*, but from the mismatch it creates between rostral neurons whose *Stxbp1* expression has been normalized and caudal neurons that remain haploinsufficient. According to the Allen Mouse Brain Atlas (https://mouse.brain-map.org/experiment/show/1010) [18], *Stxbp1* is highly expressed across caudal regions including thalamus, hypothalamus, and brainstem; in *Stxbp1*^+/−^ mice, these populations likely remain at approximately half the normal Munc18-1 level even after CRISPR-ON treatment. Such regional imbalance may destabilize the coordinated activity of brainstem circuits that govern autonomic and respiratory functions.

This interpretation is further reinforced by the striking mortality observed when *Vglut2-Cre*, which is enriched in caudal subcortical and brainstem excitatory neurons, was co-administered with *EF1a*-Cre and the 4xgRNA cassette to extend CRISPR-ON activity across both rostral and caudal regions. In *Stxbp1*^+/−^ mice, four of five animals that received this combination died between postnatal weeks 3 and 4. Although extending CRISPR-ON coverage to caudal regions might *a priori* have been expected to ameliorate the rostral–caudal expression mismatch discussed above, the severe lethality of this treatment indicates the opposite outcome: the strong transcriptional activation achieved by CRISPR-ON in caudal *Vglut2*+ neurons appears to drive *Stxbp1* expression beyond a tolerable range, perturbing neural activity in these populations in a way that can be fatal even when the underlying genotype is *Stxbp1*^+/−^. Thus, the level of Munc18-1 in caudal brainstem and subcortical excitatory neurons appears to be tightly constrained, with acute over-induction being poorly tolerated. In addition, because the *Vglut2*-Cre driver does not target all neuronal populations in these caudal regions,

*Vglut2*-negative cells, including inhibitory neurons, would be expected to remain haploinsufficient. The resulting imbalance in *Stxbp1* expression, with elevated levels in *Vglut2*+ excitatory neurons but persistently reduced levels in neighboring *Vglut2*-negative cells, may further disrupt the coordinated activity of caudal circuits and thereby contribute to the increased risk of sudden death. More specifically, these findings raise the possibility that, within the caudal brain regions targeted by *Vglut2*-Cre, particular neural circuits exist in which elevation of *Stxbp1* expression disrupts the balance of neural activity with lethal consequences. Taken together, these observations have an important translational implication: for *Stxbp1* gene activation therapy, uniform restoration of expression to near-physiological levels across the brain, rather than regional rescue or unrestrained activation in caudal populations, is likely to be the most desirable outcome, and careful titration of activation magnitude will be essential to avoid overexpression toxicity in vulnerable caudal populations. Whether elevated *STXBP1* expression is tolerated in humans is not known; no clinical phenotype has been firmly attributed to increased *STXBP1* dosage, and this uncertainty is an important consideration for translation. Although the toxicity observed here represents an unexpected finding, it provides valuable guidance for the development of gene activation therapies, highlighting the importance of regulating both the spatial distribution and the magnitude of transcriptional activation. Such considerations will be critical for the safe design of CRISPR-ON–based and other gene activation strategies for *STXBP1*-related and other haploinsufficiency disorders. This interpretation remains a hypothesis. Excessive activation in particular neuronal populations, activation in non-neuronal cells, and off-target transcriptional effects of dCas9-VPR are not excluded by the present data, and distinguishing among these possibilities will require longitudinal monitoring with time-course expression analysis, together with a comparison of survival across vector doses and gRNA numbers.

The dCas9-VPR transcriptional activator has previously been associated with cellular toxicity attributable to the VPR domain [16]. In the present study, wild-type mice expressing dCas9-VPR in the brain in the absence of *Stxbp1*-targeting gRNAs exhibited hyperactivity in the open field test, including increased locomotion and center-zone occupancy, together with a trend toward increased aggression in the resident-intruder test. These dCas9-VPR–associated changes were not detected in CRISPR-ON–treated *Stxbp1*^+/−^ mice, whose open-field measures were comparable to those of untreated *Stxbp1*^+/−^ controls. Because dCas9-VPR expression alone tended to increase rather than decrease aggression, it is unlikely to have produced the lower attack counts observed in CRISPR-ON–treated mice; however, given the small group sizes, a contribution of dCas9-VPR expression to the phenotypic measures reported here cannot be excluded. Expression of dCas9-VPR requires Cre, so the untreated *Stxbp1*^+/−^ controls, which received saline, did not express it; treated and untreated *Stxbp1*^+/−^ groups therefore differ both in dCas9-VPR expression and in *Stxbp1* activation. A *Stxbp1*^+/−^ group receiving Cre without *Stxbp1*-targeting gRNAs was not included, so these two contributions cannot be separated in the present design. Future iterations of this approach will benefit from engineering more compact, lower-toxicity transcriptional activators and from gRNA designs that maximize on-target specificity.

Limitations. This study has several limitations. First, the in vivo phenotypic analyses were performed in small cohorts: electrocorticographic recordings were obtained from only two animals per *Stxbp1*^+/−^ group, and three CRISPR-ON–treated *Stxbp1*^+/−^ mice were available for behavioral testing. These cohorts provide no statistical power to detect phenotypic rescue, and the apparent trends toward lower SWD frequency and fewer attacks must be regarded as descriptive. Second, CRISPR-ON treatment reduced survival for reasons that remain undetermined, which is itself a concern for translation; generating larger cohorts would therefore entail the loss of additional animals and could not be justified under the 3Rs principle of Reduction; adequately powered phenotypic studies will therefore require prior optimization of the vector system to eliminate this mortality. Third, the spatial and cellular distribution of CRISPR-ON activity was inferred from a Cre reporter line rather than measured directly in treated animals, so the proportion and identity of neurons in which *Stxbp1* was activated remain undefined. Fourth, transcriptome-wide analyses were not performed, and off-target transcriptional effects of dCas9-VPR cannot be excluded. Fifth, the in vitro gRNA screen was performed with three independent transfections per guide, and only gRNAs #11 and #12 were statistically distinguishable from the control conditions, so the rank order of the individual gRNAs should be regarded as exploratory rather than as a quantitative measure of relative potency. Because transcript levels were measured in bulk transfected populations without isolation of transfected cells, the activation reported here also underestimates the per-cell magnitude of activation, and differences in transfection efficiency between wells are the most likely source of the variability among replicates. Addressing these limitations is the central objective of our ongoing work.

In summary, CRISPR-ON-mediated activation of endogenous *Stxbp1* raised brain *Stxbp1* mRNA above, and restored Munc18-1 protein to, wild-type levels in *Stxbp1*^+/−^ mice. Whether this molecular correction translates into phenotypic benefit could not be determined from the present cohorts: SWD frequency and attack counts tended to be lower in treated mice, but neither comparison was statistically supported, reduced body weight was not rescued, and survival was decreased. Unlike conventional CRISPR gene editing, CRISPR-ON avoids permanent genomic modification. Together with our earlier work in *Scn1a*-haploinsufficient mice [16], the present findings establish that endogenous gene activation can normalize the protein level of a haploinsufficient gene in the brain *in vivo*, while demonstrating that uniform, appropriately titrated activation across brain regions will be required before therapeutic efficacy can be assessed.

## Materials and methods

### Mice

All mice were maintained on a 12 h light/dark cycle with ad libitum access to food and water. To generate *Stxbp1*^+/−^ mice, *Stxbp1*^fl/+^ mice (RIKEN BRC: RBRC05604) were crossed with *EIIA*-Cre transgenic mice, which express Cre recombinase ubiquitously (kindly provided by Dr. Shigemi Itohara, RIKEN Center for Brain Science), to excise exon 3 of *Stxbp1* and generate *Stxbp1*^+/−^/*EIIA*-Cre offspring. These mice were subsequently crossed with C57BL/6N mice (Jackson Laboratory Japan), and Cre-negative progeny were used as *Stxbp1*^+/−^ mice [6]. The line was thereafter maintained by continued backcrossing of *Stxbp1*^+/−^ mice to C57BL/6N. To obtain *Stxbp1*^+/+^/dCas9-VPR^fl/+^ and *Stxbp1*^+/−^/dCas9-VPR^fl/+^ mice, flox-dCas9-VPR KI-Tg mice [16], which carry a Cre-inducible Sp-dCas9-VPR cassette knocked into the mouse *Rosa26* locus, were crossed with *Stxbp1*^+/−^ mice by either natural mating or in vitro fertilization followed by embryo transfer; the flox-dCas9-VPR KI-Tg line was maintained by backcrossing to C57BL/6J mice (Jackson Laboratory Japan). To visualize Cre-mediated recombination, B6.Cg-Gt(ROSA)26Sortm14(CAG-tdTomato)Hze/J (ROSA-tdTomato; Jackson Laboratory, JAX:007914) mice were used. Genotyping was performed by PCR using the following primer pairs: *Stxbp1* (5′-GGTGGTGGACCAGTTAAGCATG-3′, 5′-TGGCAGCAAACACTATCGATAAG-3′, and 5′-CCAGCCCCAGTCTTCTTTCTC-3′; floxed and wild-type alleles, 288 bp; knockout allele, 186 bp); Cre (5′-AGGTTCGTTCACTCATGGA-3′ and 5′-TCGACCAGTTTAGTTACCC-3′; Cre+, 235 bp); and dCas9-VPR (5′-GTATCTGGCCAGCCACTATG-3′ and 5′-TGATATCAACGCGTCAAGTCG-3′; dCas9−, 0.3 kb; dCas9+, 0.5 kb).

### Designing of the gRNA sequences

The GRCm38.p6 reference sequence of the C57BL/6J strain (Ensembl, https://asia.ensembl.org/Mus_musculus/Info/Annotation) and the GRCh37/hg19 reference sequence (UCSC Genome Browser, http://genome.ucsc.edu/cgi-bin/hgGateway) were used for mouse and human, respectively. To identify candidate target sites for transcriptional activation, the ∼1 kb region immediately upstream of the transcription start site (TSS) of *Stxbp1* was compared between mouse and human, and a region exhibiting high sequence conservation between the two species was selected for gRNA design. Within this conserved region, fourteen 20-nucleotide gRNA target sequences, each immediately followed by an NGG protospacer-adjacent motif (PAM) compatible with *Streptococcus pyogenes* Cas9 (SpCas9), were selected to tile across the promoter. To avoid off-target effects, the targeting sequences were selected so as not to be identical or highly homologous to other genomic regions, as assessed by BLASTN search. The resulting fourteen gRNAs were used to construct the individual-gRNA plasmids described below (see Expression constructs).

### Expression constructs

All constructs were generated using standard molecular cloning techniques, and the insert region of every final construct was verified by Sanger sequencing. Detailed cloning procedures and all primer sequences are provided in the Supplementary Materials (**Supplementary Methods** and **Supplementary Tables S1–S3**).

Fourteen individual-gRNA constructs targeting a region of the *Stxbp1* promoter conserved between human and mouse were generated by cloning annealed oligonucleotide pairs into MLM3636 (a gift from Dr. Keith Joung; Addgene #43860). Based on *in vitro* screening, the most active gRNAs were assembled into multiplexed 3xgRNA (#10, #11, #12) and 4xgRNA (#4, #10, #11, #12) cassettes. These cassettes were subcloned into AAV backbones derived from pAAV-MCS (Agilent) to generate Cre-expressing constructs under the *EF1a* core or *CMV* promoter (pAAV-*EF1a* core-Cre and pAAV-*CMV*-Cre), the corresponding Cre-gRNA dual-expression constructs (e.g., pAAV-*EF1a* core-Cre-4x*Stxbp1* gRNA), and gRNA-only constructs (pAAV-3x/4x*Stxbp1* gRNA). For *in vitro* evaluation, Sp-dCas9-VPR (a gift from Dr. George Church; Addgene #63798) and a Sp-dCas9-VPR-IRES-EGFP construct were used. pAAV-*Vglut2*-mCherry-P2A-Cre was generated by assembling a pAAV backbone, a mouse *Vglut2* promoter fragment, and an mCherry-P2A-Cre cassette. The AAV-PHP.eB capsid plasmid (pUCmini-iCAP-PHP.eB; a gift from Dr. Viviana Gradinaru; Addgene #103005) was used.

### Evaluation of gRNAs in cell culture

Neuro2A cells were seeded in 6-well plates at 1 × 10⁵ cells per well two days prior to transfection and maintained in Dulbecco’s Modified Eagle Medium (DMEM; Thermo Fisher Scientific) supplemented with 10% fetal bovine serum (FBS; GIBCO) at 37 °C in a humidified incubator with 5% CO₂. When cells reached approximately 80% confluence, they were transfected with plasmid DNA using Lipofectamine LTX Reagent with PLUS (Thermo Fisher Scientific). For evaluation of individual gRNAs, 500 ng of Sp-dCas9-VPR plasmid was co-transfected with 500 ng of a single gRNA plasmid per well. For evaluation of multiplexed gRNAs (combinations of two to four gRNAs), 1 µg of Sp-dCas9-VPR plasmid was co-transfected with a total of 500 ng of gRNA plasmid mixture per well. At 48 h post-transfection, total RNA was extracted from the transfected cells using TRIzol Reagent (Thermo Fisher Scientific).

### Real-time quantitative PCR (qPCR) analysis

Total RNA was extracted from cultured Neuro2A cells or from mouse brain hemispheres collected at 5 weeks of age, using TRIzol Reagent (Thermo Fisher Scientific). One microgram of total RNA was reverse-transcribed into complementary DNA (cDNA) using the PrimeScript RT Reagent Kit with gDNA Eraser (Takara Bio) according to the manufacturer’s instructions. Quantitative PCR was performed on a 7900HT Fast Real-Time PCR System (Thermo Fisher Scientific) using SsoAdvanced Universal SYBR Green Supermix (Bio-Rad), with *Gusb* (β-glucuronidase) as the housekeeping reference gene. The following primers were used: *Stxbp1*, 5′-TCACCAACATGGCTCACCTC-3′ (forward) and 5′-AATGTCCATAGCGGGCACTC-3′ (reverse); *Gusb*, 5′-ACTGACACCTCCATGTATCCCAAG-3′ (forward) and 5′-CAGTAGGTCACCAGCCCGATG-3′ (reverse). Each sample was assayed in triplicate, and relative *Stxbp1* expression was calculated using the ΔΔCt method. For *in vitro* experiments, untransfected Neuro2A cells or cells transfected with an empty vector (MLM3636 lacking a target gRNA sequence) served as controls. Transfection efficiency was not measured in individual wells, and transfected cells were not enriched by sorting or identified by a co-transfected reporter; *Stxbp1* mRNA was therefore quantified in the whole transfected population.

### Western blot analysis

Neuro2A cells (cultured in 12-well plates) or mouse brain hemispheres collected at 5 weeks of age were lysed in RIPA buffer (150 mM NaCl, 50 mM Tris-HCl pH 7.5, 1% NP-40, 0.5% sodium deoxycholate, 0.1% SDS) supplemented with a protease inhibitor cocktail (cOmplete; Sigma-Aldrich), using 0.5 mL per well of cells or 5 mL per brain hemisphere. Samples were homogenized by pipetting, glass Dounce homogenization, or hand-held sonication, and then centrifuged at 12,000 rpm for 10 min at 4 °C. The supernatants were collected, and 15 µg of total protein per sample was separated by SDS-PAGE on a SuperSep Ace 5–20% gradient gel (13-well; Wako). Proteins were transferred onto a PVDF membrane (Bio-Rad), which was then blocked in blocking buffer (1× TBS containing 0.05% Tween-20 and 5% skim milk). After washing with 1× TBS, the membrane was incubated overnight at 4 °C with primary antibody diluted in blocking buffer: anti–Munc18-1 (1:1,000; Synaptic Systems) or anti–β-tubulin (clone D66, 1:5,000; Sigma-Aldrich). Following washes with 1× TBS, the membrane was incubated for 2 h at room temperature with HRP-conjugated secondary antibody diluted in blocking buffer: donkey anti-rabbit IgG (H+L)–HRP (1:10,000; Jackson ImmunoResearch) or anti-mouse IgG (H+L)–HRP (1:5,000; Promega), respectively. Immunoreactive bands were visualized using Western Lightning Plus ECL (PerkinElmer) and imaged on an Amersham Imager 600 (GE Healthcare).

### Histological analyses

Mice were deeply anesthetized with a combination of medetomidine (0.3 mg/kg), midazolam (4 mg/kg), and butorphanol (5 mg/kg) administered intraperitoneally, and transcardially perfused with 0.9% saline, followed by 4% paraformaldehyde in 0.1 M phosphate buffer (pH 7.4). Brains were removed and embedded in Tissue-Tek O.C.T. compound (Sakura Finetek Japan). Sagittal sections (30 µm thick) were prepared using a CM3050 cryostat (Leica Biosystems). tdTomato fluorescence images were acquired using a BZ-X710 fluorescence microscope (Keyence).

### AAVs

AAV vectors were packaged into the AAV-PHP.eB capsid as previously described [16, 19, 20]. The following plasmids were packaged into AAV-PHP.eB particles for *in vivo* delivery: pAAV-*EF1a* core-Cre-4x*Stxbp1* gRNA, pAAV-*EF1a* core-Cre (Cre-only control), and pAAV-*Vglut2*-mCherry-P2A-Cre. The resulting viral particles are referred to as AAV-PHP.eB-*EF1a* core-Cre-4x*Stxbp1* gRNA, AAV-PHP.eB-*EF1*a core-Cre, and AAV-PHP.eB-*Vglut2*-mCherry-P2A-Cre, respectively.

### AAV administration

#### Retro-orbital injection

AAV-PHP.eB-*Vglut2*-mCherry-P2A-Cre (4.5 × 10¹⁰ vg/mouse) was administered into the right retro-orbital sinus of ROSA-tdTomato mice (10 weeks old), as previously described [21]. The virus was diluted in sterile saline to a final volume of 150 µL per mouse and delivered using an insulin syringe with an integrated needle (MyJector; Terumo). Injected mice were used for tdTomato fluorescence imaging at 15 weeks of age.

#### Intracerebroventricular (ICV) injection

Neonatal (postnatal day 1) *Stxbp1*^+/+^/dCas9-VPR^fl/+^ and *Stxbp1*^+/−^/dCas9-VPR^fl/+^ mice were anesthetized by hypothermia on an aluminum block chilled on ice for 4 min, following a previously described procedure [22]. AAV-PHP.eB-*EF1a*-Cre-4x*Stxbp1* gRNA (5.7 × 10¹⁰ vg/mouse), AAV-PHP.eB-*EF1a*-Cre (5.6 × 10¹⁰ vg/mouse), or an equivalent volume of sterile saline was then bilaterally injected into the lateral ventricles using a 30 G × 15 mm Type-A microsyringe-compatible needle (ITO CORPORATION) inserted perpendicular to the skull to a depth of 2 mm. Each injection site was defined as the point that internally divides the line connecting lambda and the ipsilateral eye in a 2:5 ratio (from lambda). Viruses were diluted in sterile saline, and all injections were delivered at 2 µL per hemisphere (4 µL/mouse total). All injection solutions contained 0.05% Trypan Blue Solution (Gibco) to allow visual confirmation of successful delivery.

### Electrocorticogram (ECoG) recording

Mice were anesthetized with isoflurane (1.0–2.5%) and placed in a stereotaxic frame (Stoelting). After disinfecting the scalp with chlorhexidine, a midline incision was made to expose the skull. Holes were drilled with a hand drill over the bilateral somatosensory cortex (coordinates relative to bregma: AP −1.00 mm, ML ±1.50 mm), and screw electrodes were inserted for ECoG recording. A pair of wire electrodes was implanted into the dorsal neck muscles for electromyography (EMG). The electrodes were fixed in place with dental cement, and mice were allowed to recover for at least one week before recording. ECoG and EMG signals were recorded continuously for approximately 48 h with mice freely moving and with *ad libitum* access to food and water. The frequency of spike-wave discharges (SWDs) was quantified over a 24-h window (08:00 to 08:00 the following day) on the second day of recording. SWDs were defined, as described previously [12], as bursts of negative deflections at 3–10 Hz with amplitudes exceeding two-fold the baseline amplitude and comprising at least three consecutive cycles. All analyses were performed by an investigator blinded to mouse genotype.

### Resident-intruder test

Aggressive behavior was assessed at 16 weeks of age using the resident-intruder test. From 5 weeks of age, male resident test mice were singly housed in standard cages (182 × 260 × 128 mm), and the bedding was left unchanged during the week preceding the test to promote territorial behavior. On the test day, a male C57BL/6NCrSlc mouse (6–7 weeks old; Japan SLC) was introduced into the resident’s home cage as an intruder, and the encounter was video-recorded from above under approximately 70 lux illumination for 10 min. Offline video analysis was performed by an investigator blinded to the resident’s genotype, who scored the frequency of aggressive behaviors (attacks and chasing) and non-aggressive behaviors (mounting and sniffing).

### Open field test

Male mice were habituated to the test room for at least 30 min prior to testing. Each mouse was then placed in a corner of a square open-field apparatus (45 × 45 × 40 cm; PanLab) illuminated at 100 lx and allowed to freely explore for 30 min. The session was video-recorded, and the percentage of time spent in the center area (20 × 20 cm), total distance traveled (cm), and locomotor velocity (cm/s) were automatically scored using the manufacturer’s software (Smart 3.0; PanLab). Between trials, the apparatus was cleaned with 70% ethanol.

### Statistical analyses

All statistical analyses were performed using KyPlot 6.0 (KyensLab). Comparisons among three or more groups were performed using one-way analysis of variance (ANOVA), followed by Tukey’s post-hoc test for pairwise comparisons among all groups or Dunnett’s post-hoc test for comparisons against a single control group. Before parametric testing, normality was assessed by the Shapiro-Wilk test applied to the residuals of each ANOVA model, and homogeneity of variance was assessed by Levene’s test with median centring; these tests pool information across groups, which is necessary because the per-group sample size is too small for group-wise assessment.

Neither assumption was violated in any of the datasets analyzed parametrically, and the results are reported in **Supplementary Table S6**. The attack counts in Figure 4C are discrete and markedly skewed, and the group sizes are unequal, so these data were compared by the Kruskal-Wallis test with post-hoc comparisons rather than by ANOVA. Deaths during the rearing period were ascertained at intermittent cage checks, so the exact time of death is not known for individual animals; survival is therefore presented as the proportion of animals alive at the end of each 5-day interval, and no time-to-event analysis was performed. Groups were compared by Fisher’s exact test on survival at postnatal day 35. Data are presented as the mean ± standard error of the mean (SEM). Statistical significance was defined as *p* < 0.05 (\**p* < 0.05, \*\**p* < 0.01, \*\*\**p* < 0.001).

## Acknowledgments

The authors thank the members of the Section of Viral Vector Development at National Institute for Physiological Sciences, the Department of Neurodevelopmental Disorder Genetics and the animal facility at Nagoya City University (NCU) for their support. We are grateful to the Research Equipment Sharing Center at NCU for technical assistance. We also thank Ms. Nana Nagata for her contribution to part of the evaluation of gRNA efficacy in vitro in the early phase of this study.

## Author Contributions

TY, KY and TS contributed to the study conception and design. Material preparation, data collection, and analysis were performed by TY, RI, KK, MS, YH, KY and TS. The first draft of the manuscript was written by TY, RI, KY and TS. All authors have reviewed and approved the final manuscript.

## Data Availability

The datasets generated during and/or analyzed during the current study are available from the corresponding author on reasonable request.

## Ethics Approval

All animal breeding and experimental procedures were approved by the Institutional Animal Care and Use Committee of NCU (approval No. 19-032, approved 20 Dec 2022; approval No. 23-024, approved 21 Apr 2023). All procedures were conducted in accordance with the ARRIVE guidelines and the institutional guidelines and regulations of NCU. This animal study adheres to the Guide for the Care and Use of Laboratory Animals.

## Consent to Participate

Not applicable.

## Consent for Publication

Not applicable.

## Clinical trial number

Not applicable.

