## Supplementary information for "CRISPR-mediated *Stxbp1* gene activation restores Munc18-1 levels in *Stxbp1*-haploinsufficient mice"

### Supplementary Methods

#### Expression constructs

All cloning steps utilized standard molecular biology techniques. The insert region of every final construct was verified by Sanger sequencing prior to use. All primers used for cloning are listed in Supplementary Tables S1–S3.

**Individual-gRNA constructs (MLM3636-*Stxbp1* gRNA).** Fourteen candidate gRNA sequences were designed against a region of the *Stxbp1* promoter that is highly conserved between human and mouse. Complementary oligonucleotide pairs (**Supplementary Table S1**) were annealed for each gRNA and ligated into BsmBI (New England Biolabs; NEB)–digested MLM3636 (a gift from Dr. Keith Joung; Addgene #43860).

**Multiplex gRNA constructs (pCR4-3x*Stxbp1* gRNA and pCR4-4x*Stxbp1* gRNA).** Multiplex gRNA cassettes were assembled by PCR-amplifying #10, #11, and #12 (for 3x*Stxbp1* gRNA) or #4, #10, #11, and #12 (for 4x*Stxbp1* gRNA) (primers in **Supplementary Table S2)** from the corresponding MLM3636-*Stxbp1* gRNA templates. Amplicons were digested with BbsI (NEB) and ligated, and the resulting ~1.5 kb (3xgRNA) or ~2.0 kb (4xgRNA) products were cloned into pCR4-Blunt-TOPO (Invitrogen).

**pAAV-*EF1a* core-Cre.** The *EF1a* core promoter, Cre-WPREγ, and WPRE stem-loop fragments were PCR-amplified from pAAV-*EF1a*-mCherry-IRES-Cre (Addgene #55632), and the SPA sequence was generated by primer-only PCR (all primers in **Supplementary Table S3**). These fragments were assembled with a 2.9-kb pAAV-MCS (Agilent) backbone (prepared by MluI/PmlI (NEB) digestion) using the In-Fusion HD Cloning Kit (Takara Bio).

**pAAV-CMV-Cre.** A Cre-WPRE cassette amplified from pAAV-*EF1a*-mCherry-IRES-Cre (primers in **Supplementary Table S3**) was assembled with a 5.7-kb pAAV-MCS backbone (prepared by ClaI/BglII [NEB] digestion) using the In-Fusion HD Cloning Kit.

**Cre-gRNA dual-expression AAV constructs.** To generate pAAV-*EF1a core-Cre*-3x*Stxbp1* gRNA, pAAV-*EF1a core-Cre*-4x*Stxbp1* gRNA, pAAV-*CMV-Cre*-3x*Stxbp1* gRNA, and pAAV-*CMV-Cre*-4x*Stxbp1* gRNA, the 3x*Stxbp1* gRNA or 4x*Stxbp1* gRNA cassette was excised from the corresponding pCR4 plasmid with EcoRI (NEB) and ligated into the appropriate Cre-expression AAV backbone (pAAV-*EF1a core-Cre* or pAAV-*CMV-Cre*).

**gRNA-only AAV constructs (pAAV-3x*Stxbp1* gRNA and pAAV-4x*Stxbp1* gRNA).** The 2.9-kb pAAV-MCS backbone (NotI [NEB]–digested) was ligated with the 3x*Stxbp1* gRNA or 4x*Stxbp1* gRNA cassette.

**Sp-dCas9-VPR-IRES-EGFP.** Sp-dCas9-VPR (Addgene #63798) was digested with NotI (NEB) and blunted. An IRES-EGFP fragment, PCR-amplified from pIRES2-EGFP (Clontech) using primers in **Supplementary Table S3**, was inserted into this site.

**pAAV-*Vglut2*-mCherry-P2A-Cre.** A pAAV backbone (PCR-amplified from pAAV-MCS), a mouse *Vglut2* promoter fragment (amplified from C57BL/6J tail genomic DNA), and an mCherry-P2A-Cre cassette (amplified from pAAV-*EF1a*-fDIO-mCherry-P2A-Cre; Addgene #161773) were prepared using the primers listed in **Supplementary Table S3**. The three fragments were assembled using the In-Fusion HD Cloning Kit.

**pUCmini-iCAP-PHP.eB.** Plasmid DNA encoding the AAV-PHP.eB capsid was generously provided by Dr. Viviana Gradinaru (California Institute of Technology).

**Supplementary Table S1. Oligonucleotide sequences for individual-gRNA construct generation.**

| gRNA No. | Forward primer | Reverse primer |
|---|---|---|
| *Stxbp1* gRNA #1 | acaccggaatgctaagactttaactgg | aaaaccagttaaagtcttagcattccg |
| *Stxbp1* gRNA #2 | acaccgttattttattttccaccatcg | aaaacgatggtggaaaataaaataacg |
| *Stxbp1* gRNA #3 | acaccgaagcaagtgctccacctctgg | aaaaccagaggtggagcacttgcttcg |
| *Stxbp1* gRNA #4 | acaccgcagtatgtgaccatgtagacg | aaaacgtctacatggtcacatactgcg |
| *Stxbp1* gRNA #5 | acaccgtcctaattagatttggacttg | aaaacaagtccaaatctaattaggacg |
| *Stxbp1* gRNA #6 | acaccgacatagcttatacatagttag | aaaactaactatgtataagctatgtcg |
| *Stxbp1* gRNA #7 | acaccgggtacaacagcaagcgcctcg | aaaacgaggcgcttgctgttgtacccg |
| *Stxbp1* gRNA #8 | acaccgagagcttagctctccatcaag | aaaacttgatggagagctaagctctcg |
| *Stxbp1* gRNA #9 | acaccgcctatgttgtggctgggaatg | aaaacattcccagccacaacataggcg |
| *Stxbp1* gRNA #10 | acaccgtggaaggctcccaaaaaagcg | aaaacgcttttttgggagccttccacg |
| *Stxbp1* gRNA #11 | acaccgatgacctgggcagcccgtgcg | aaaacgcacgggctgcccaggtcatcg |
| *Stxbp1* gRNA #12 | acaccgtggcgcgagcgtgaggcgcag | aaaactgcgcctcacgctcgcgccacg |
| *Stxbp1* gRNA #13 | acaccgcgcccgggccgcgcttctatg | aaaacatagaagcgcggcccgggcgcg |
| *Stxbp1* gRNA #14 | acaccgcccgcccctcccgcgcgcgcg | aaaacgcgcgcgcgggaggggcgggcg |

**Supplementary Table S2. Primer sequences for multiplexed (3x and 4x) gRNA construct generation.**

| | Template | Forward primer | Reverse primer |
|---|---|---|---|
| 3xgRNA | MLM3636-*Stxbp1*gRNA #10 | gatcggatccggtaccaaggt | cgatcgaagacttgtgttcctggccttttgctggcct |
| | MLM3636-*Stxbp1*gRNA #11 | cgatcgaagacttacacgatcggatccggtaccaaggt | cgatcgaagacttcacgtcctggccttttgctggcct |
| | MLM3636-*Stxbp1*gRNA #12 | cgatcgaagacttcgtggatcggatccggtaccaaggt | tcctggccttttgctggcct |
| 4xgRNA | MLM3636-*Stxbp1*gRNA #4 | gatcggatccggtaccaaggt | cgatcgaagacttgtgttcctggccttttgctggcct |
| | MLM3636-*Stxbp1*gRNA #10 | cgatcgaagacttacacgatcggatccggtaccaaggt | cgatcgaagacttcacgtcctggccttttgctggcct |
| | MLM3636-*Stxbp1*gRNA #11 | cgatcgaagacttcgtggatcggatccggtaccaaggt | cgatcgaagacttcctttcctggccttttgctggcct |
| | MLM3636-*Stxbp1*gRNA #12 | cgatcgaagacttaagggatcggatccggtaccaaggt | tcctggccttttgctggcct |

**Supplementary Table S3. Primer sequences for expression construct generation.**

| | Forward primer | Reverse primer |
|---|---|---|
| *EF1a* core promoter | ctgcggccgcacgcgggctccggtgcccgtcagtg | tgtggccatggtggcctgtgttctggcggcaaacc |
| Cre-WPREγ | gccaccatggccacaaccatgcccaag | gaactaaccaggatttatacaaggaggag |
| WPRE stem loop region | aatcctggttagttcttgccacggcggaactcatc | ccacggaattgtcagtgcccaac |
| SPA sequence | ctgacaattccgtggaataaaagatctttattttcattagatctgtgtgttggttttttgtgtg | ccgctcggtccgcacgtgcacacaaaaaaccaacacacagatctaatgaaaataaagatcttttatt |
| Cre-WPRE sequence | gggattcgaacatcgatgccaccatggccacaaccatgc | gccacccgtagatctgcggggaggcggcccaaagg |
| IRES-EGFP | ggatccgcccctctccct | gctttacttgtacagctcgtcc |
| pAAV backbone | attcgatatcaagcttatcgataatcaacctctgg | actttatccatctttgcaaagcttacgc |
| *Vglut2* promoter fragment | aaagatggataaagtacgcactcccctggttgatttag | catggatccggtacctcttgtaaagactggtgtccagcct |
| mCherry-P2A-Cre cassette | ggtaccggatccatggtgagcaagggcgag | agcttgatatcgaatctaatcgccatcttccagcagg |

**Supplementary Table S4. Statistical comparisons for the in vitro gRNA screening experiments.**

All comparisons were made by one-way analysis of variance followed by Dunnett's post-hoc test, performed separately with each control group designated as the reference. t denotes the Dunnett test statistic for the comparison between the indicated group and the control. * P < 0.05, ** P < 0.01, *** P < 0.001.

**(A) Figure 1B. Individual gRNAs (n = 3 per group).**

| | | **Dunnett vs Neuro2A** | | **Dunnett vs NC** | |
|---|---|---|---|---|---|
| **Group** | **Mean ± SEM** | **t** | **P** | **t** | **P** |
| gRNA #1 | 0.95 ± 0.28 | -0.348 | 1.0000 | -0.180 | 1.0000 |
| gRNA #2 | 0.88 ± 0.04 | -0.584 | 0.9998 | -0.415 | 1.0000 |
| gRNA #3 | 1.20 ± 0.05 | 0.494 | 1.0000 | 0.662 | 0.9991 |
| gRNA #4 | 1.37 ± 0.26 | 1.077 | 0.9442 | 1.246 | 0.8705 |
| gRNA #5 | 0.79 ± 0.09 | -0.864 | 0.9891 | -0.696 | 0.9985 |
| gRNA #6 | 1.09 ± 0.15 | 0.146 | 1.0000 | 0.314 | 1.0000 |
| gRNA #7 | 1.20 ± 0.22 | 0.494 | 1.0000 | 0.662 | 0.9991 |
| gRNA #8 | 1.49 ± 0.10 | 1.470 | 0.7281 | 1.638 | 0.6066 |
| gRNA #9 | 1.30 ± 0.27 | 0.853 | 0.9903 | 1.021 | 0.9610 |
| gRNA #10 | 1.45 ± 0.33 | 1.347 | 0.8112 | 1.515 | 0.6962 |
| gRNA #11 | 1.95 ± 0.31 | 3.030 | 0.0474 * | 3.198 | 0.0319 * |
| gRNA #12 | 2.39 ± 0.41 | 4.511 | 0.0010 ** | 4.679 | 0.0006 *** |
| gRNA #13 | 1.05 ± 0.11 | -0.011 | 1.0000 | 0.157 | 1.0000 |
| gRNA #14 | 0.92 ± 0.08 | -0.426 | 1.0000 | -0.258 | 1.0000 |
| NC | 1.00 ± 0.06 | -0.168 | 1.0000 | - | (control) |
| Neuro2A | 1.05 ± 0.05 | - | (control) | 0.168 | 1.0000 |

One-way ANOVA: F(15, 32) = 3.96, P = 0.0005. Neuro2A, untransfected cells; NC, cells transfected with the empty MLM3636 vector. Values are expressed relative to untransfected Neuro2A cells.

**(B) Figure 1C. Multiplexed gRNA pools (n = 3 per group).**

| | | **Dunnett vs Neuro2A** | | **Dunnett vs NC** | |
|---|---|---|---|---|---|
| **Group** | **Mean ± SEM** | **t** | **P** | **t** | **P** |
| 2xgRNA mix | 6.29 ± 0.64 | 7.397 | <0.001 *** | 7.532 | <0.001 *** |
| 3xgRNA mix | 7.24 ± 0.79 | 8.730 | <0.001 *** | 8.866 | <0.001 *** |
| 4xgRNA (a) mix | 7.26 ± 0.40 | 8.749 | <0.001 *** | 8.884 | <0.001 *** |
| 4xgRNA (b) mix | 7.04 ± 0.69 | 8.446 | <0.001 *** | 8.581 | <0.001 *** |
| 5xgRNA mix | 6.90 ± 0.32 | 8.245 | <0.001 *** | 8.385 | <0.001 *** |
| NC | 0.91 ± 0.12 | -0.135 | 1.0000 | - | (control) |
| Neuro2A | 1.00 ± 0.00 | - | (control) | 0.135 | 1.0000 |

One-way ANOVA: F(6, 14) = 33.86, P < 0.001. gRNA pools: 2xgRNA (#11, #12); 3xgRNA (#10, #11, #12); 4xgRNA (a) (#4, #10, #11, #12); 4xgRNA (b) (#8, #10, #11, #12); 5xgRNA (#4, #8, #10, #11, #12).

**(C) Supplementary Figure S1. AAV construct combinations (n = 3 per group).**

| | | **vs Neuro2A** | | **vs *EF1a*-Cre** | | **vs *CMV*-Cre** | |
|---|---|---|---|---|---|---|---|
| **Group** | **Mean ± SEM** | **t** | **P** | **t** | **P** | **t** | **P** |
| *EF1a*-Cre 4xgRNA | 6.04 ± 0.28 | 9.062 | <0.001 *** | 9.332 | <0.001 *** | 8.920 | <0.001 *** |
| *EF1a*-Cre 3xgRNA | 4.81 ± 0.63 | 6.838 | <0.001 *** | 7.109 | <0.001 *** | 6.696 | <0.001 *** |
| *CMV*-Cre 4xgRNA | 4.63 ± 0.28 | 6.520 | <0.001 *** | 6.791 | <0.001 *** | 6.378 | <0.001 *** |
| *CMV*-Cre 3xgRNA | 4.89 ± 0.41 | 6.985 | <0.001 *** | 7.255 | <0.001 *** | 6.843 | <0.001 *** |
| 4xgRNA | 4.00 ± 0.52 | 5.372 | <0.001 *** | 5.643 | <0.001 *** | 5.230 | <0.001 *** |
| 3xgRNA | 4.45 ± 0.62 | 6.195 | <0.001 *** | 6.465 | <0.001 *** | 6.053 | <0.001 *** |
| *EF1a*-Cre | 0.89 ± 0.04 | -0.270 | 1.0000 | - | (control) | -0.412 | 0.9992 |
| *CMV*-Cre | 1.11 ± 0.01 | 0.142 | 1.0000 | 0.412 | 0.9992 | - | (control) |
| Neuro2A | 1.04 ± 0.02 | - | (control) | 0.270 | 1.0000 | -0.142 | 1.0000 |

One-way ANOVA: F(8, 18) = 25.54, P < 0.001. Neuro2A, untransfected cells. Cells transfected with a Cre construct alone (*EF1a*-Cre or *CMV*-Cre) served as additional controls, and the analysis was repeated with each of the three control groups as the reference. All six construct-containing groups differed significantly from every control group.

**Supplementary Table S5. Statistical comparisons for brain *Stxbp1* mRNA and Munc18-1 protein levels (Figures 3A and 3B).**

All comparisons were made by one-way analysis of variance followed by Tukey's post-hoc test. P values are given for every pairwise comparison, including those that did not reach significance. * P < 0.05, ** P < 0.01, *** P < 0.001.

**(A) Figure 3A. Brain *Stxbp1* mRNA, expressed relative to untreated *Stxbp1*+/+ mice.**

| Group | n | Mean ± SEM | vs (1) | vs (2) | vs (3) | vs (4) |
|---|---|---|---|---|---|---|
| (1) *Stxbp1*+/+, Cre + 4xgRNA | 3 | 2.81 ± 0.13 | - | <0.001 *** | <0.001 *** | <0.001 *** |
| (2) *Stxbp1*+/−, Cre + 4xgRNA | 3 | 1.53 ± 0.12 | <0.001 *** | - | <0.001 *** | 0.0018 ** |
| (3) *Stxbp1*+/−, untreated | 4 | 0.54 ± 0.03 | <0.001 *** | <0.001 *** | - | 0.0035 ** |
| (4) *Stxbp1*+/+, untreated | 5 | 0.99 ± 0.03 | <0.001 *** | 0.0018 ** | 0.0035 ** | - |

One-way ANOVA: F(3, 11) = 154.72, P < 0.001. "Cre + 4xgRNA" denotes administration of AAV-PHP.eB-*EF1a*-Cre-4xgRNA; untreated animals received saline. Tissue was collected at 5 weeks of age.

**(B) Figure 3B. Brain Munc18-1 protein, expressed relative to untreated *Stxbp1*+/+ mice.**

| Group | n | Mean ± SEM | vs (1) | vs (2) | vs (3) | vs (4) |
|---|---|---|---|---|---|---|
| (1) *Stxbp1*+/+, Cre + 4xgRNA | 3 | 1.36 ± 0.05 | - | 0.3253 | <0.001 *** | 0.0186 * |
| (2) *Stxbp1*+/−, Cre + 4xgRNA | 3 | 1.19 ± 0.10 | 0.3253 | - | <0.001 *** | 0.2369 |
| (3) *Stxbp1*+/−, untreated | 3 | 0.48 ± 0.04 | <0.001 *** | <0.001 *** | - | 0.0020 ** |
| (4) *Stxbp1*+/+, untreated | 3 | 1.00 ± 0.04 | 0.0186 * | 0.2369 | 0.0020 ** | - |

One-way ANOVA: F(3, 8) = 34.81, P < 0.001. Groups and treatments are as in (A). Munc18-1 signal intensity was normalized to β-tubulin.

**Supplementary Table S6. Tests of model assumptions for the datasets analyzed by one-way analysis of variance.**

k, number of groups; N, total number of observations. Normality was assessed by the Shapiro-Wilk test applied to the residuals of each ANOVA model, which pools information across groups; homogeneity of variance was assessed by Levene's test with median centring, which is robust to departures from normality. Bartlett's test is shown for completeness. The corresponding analysis-of-variance results and post-hoc comparisons are given in Supplementary Tables S4 and S5.

| Dataset | k | N | Shapiro-Wilk (residuals) | Levene (median) | Bartlett |
|---|---|---|---|---|---|
| Figure 1B | 16 | 48 | W = 0.970, P = 0.24 | F = 0.68, P = 0.78 | $\chi^2$ = 24.45, P = 0.058 |
| Figure 1C | 7 | 21 | W = 0.926, P = 0.11 | F = 0.62, P = 0.71 | $\chi^2$ = 23.77, P < 0.001 † |
| Figure 3A | 4 | 15 | W = 0.979, P = 0.96 | F = 0.86, P = 0.49 | $\chi^2$ = 6.55, P = 0.088 |
| Figure 3B | 4 | 12 | W = 0.954, P = 0.70 | F = 0.66, P = 0.60 | $\chi^2$ = 2.32, P = 0.51 |
| Supplementary Figure S1 | 9 | 27 | W = 0.957, P = 0.31 | F = 0.74, P = 0.66 | $\chi^2$ = 28.35, P < 0.001 † |

† Bartlett's test reaches significance for these two datasets because the normalization reference group (untransfected Neuro2A cells) has near-zero variance by construction, every value being close to 1.0; Bartlett's test is sensitive both to this and to departures from normality. With the control groups excluded, Bartlett's test is not significant (Figure 1C, five gRNA pools: $\chi^2$ = 1.71, P = 0.79; Supplementary Figure S1, six construct-containing groups: $\chi^2$ = 2.05, P = 0.84). Levene's test was therefore used as the primary criterion for homogeneity of variance.

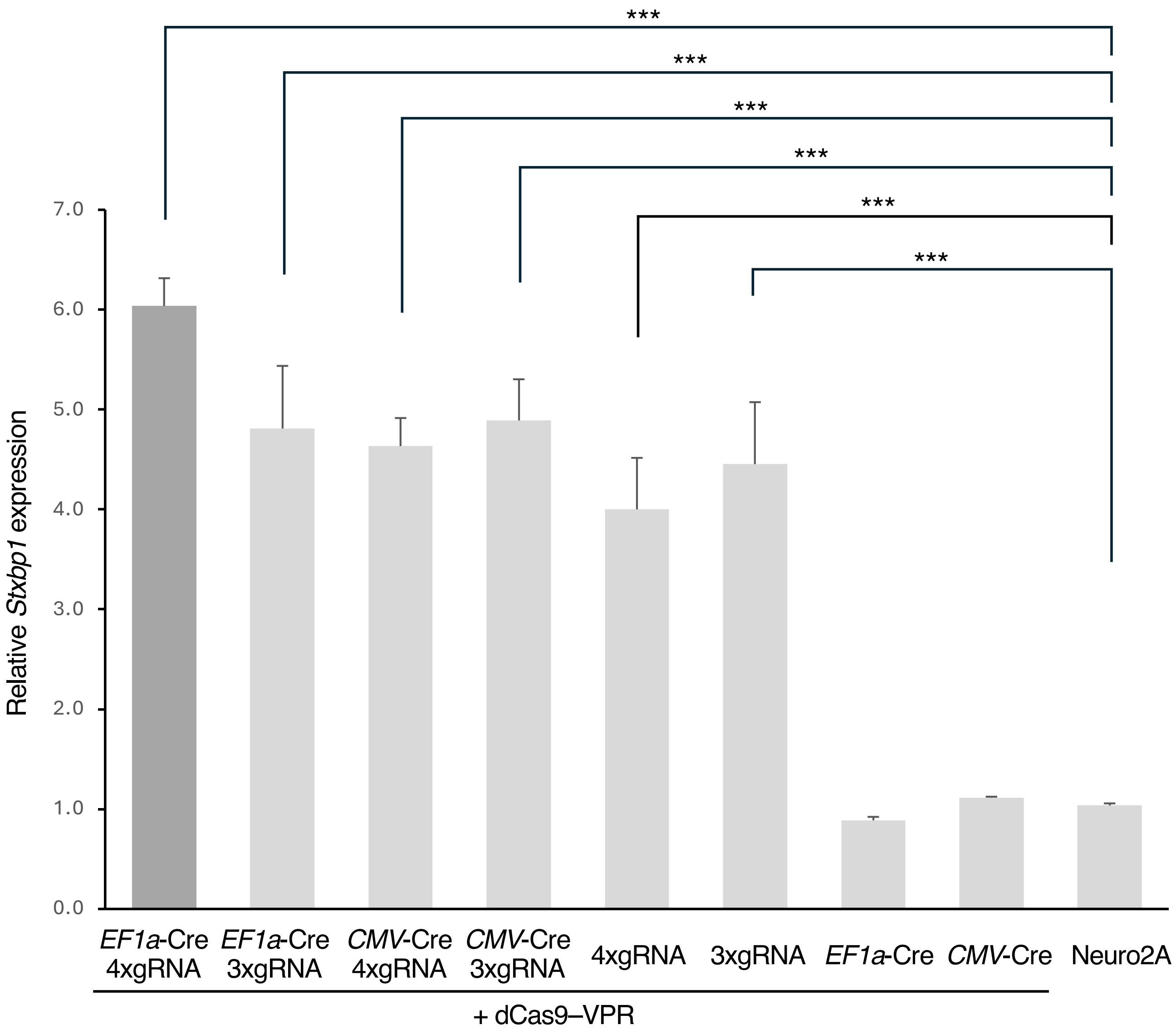


**Supplementary Figure S1. *EF1a*-Cre combined with 4xgRNA gives the highest mean transcriptional activation of *Stxbp1* among the tested Cre-driver and gRNA combinations.** Real-time qPCR quantification of *Stxbp1* mRNA levels in Neuro2A cells transfected with constructs combining a Cre driver (*EF1a*-Cre or *CMV*-Cre) and a multiplexed gRNA cassette (3xgRNA or 4xgRNA): *EF1a*-Cre 4xgRNA, *EF1a*-Cre 3xgRNA, *CMV*-Cre 4xgRNA, and *CMV*-Cre 3xgRNA. All four combination constructs increased *Stxbp1* mRNA levels 4.6- to 6.0-fold relative to control cells, with *EF1a*-Cre 4xgRNA giving the highest mean value; the differences among the construct-containing groups were not statistically significant. Cells transfected with gRNA cassettes alone (3xgRNA or 4xgRNA) or with Cre constructs alone (*EF1a*-Cre or *CMV*-Cre) were also analyzed. gRNA pools: 3xgRNA mix (#10, #11, #12); 4xgRNA mix (#4, #10, #11, #12). n = 3 per group. Statistical comparisons were performed using one-way ANOVA followed by Dunnett's post-hoc test, carried out separately with untransfected Neuro2A cells, with *EF1a*-Cre and with CMV-Cre as the reference control; asterisks denote comparisons with untransfected Neuro2A cells. All comparisons are reported in Supplementary Table S4, and the tests of model assumptions in Supplementary Table S6 (***$p < 0.001$).

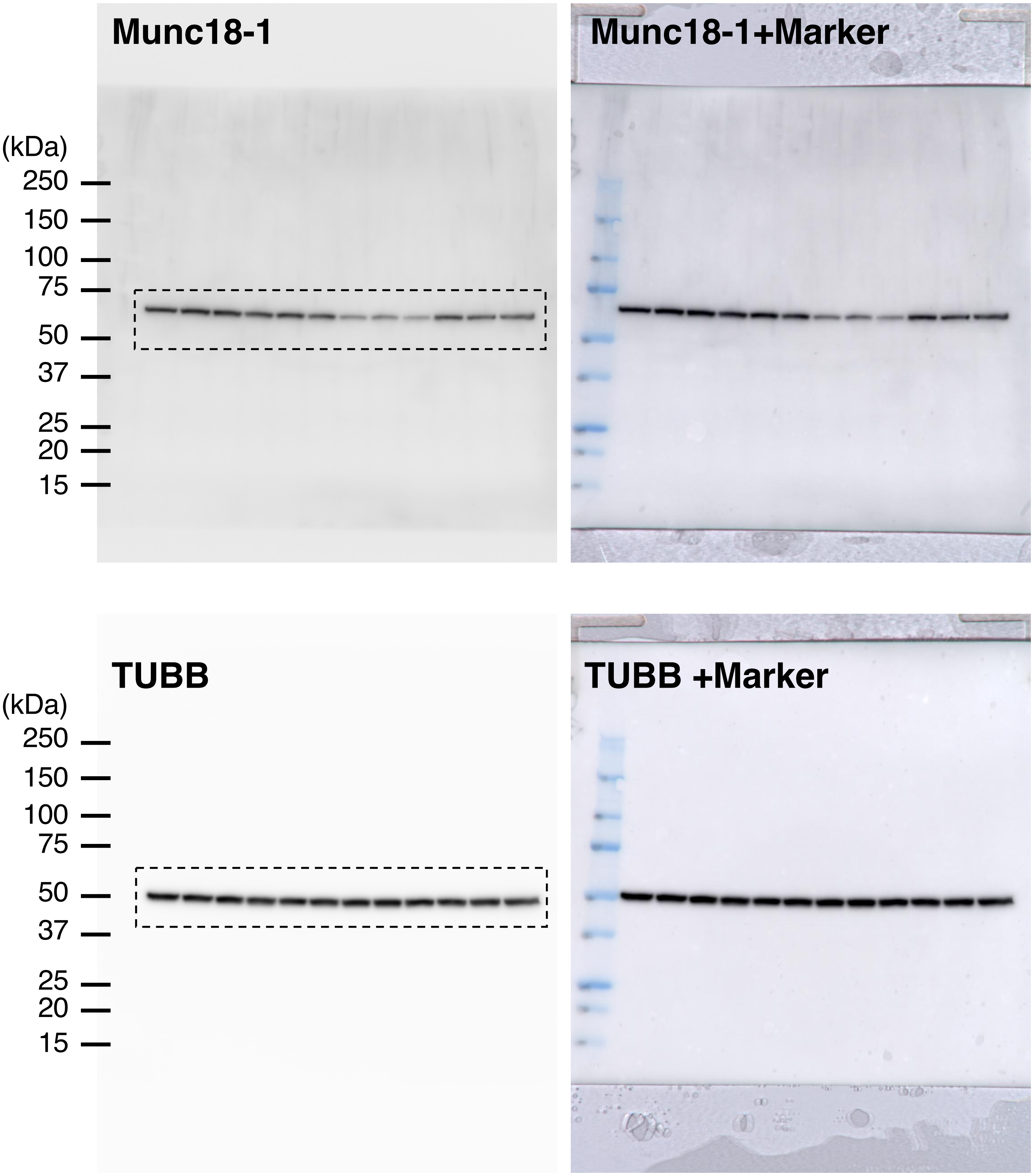


**Supplementary Figure S2. Full-size Western blot images**. Uncropped Western blot scans corresponding to the cropped panels shown in Figure 3B. Dashed rectangles indicate the regions that were cropped for presentation in the main figure.

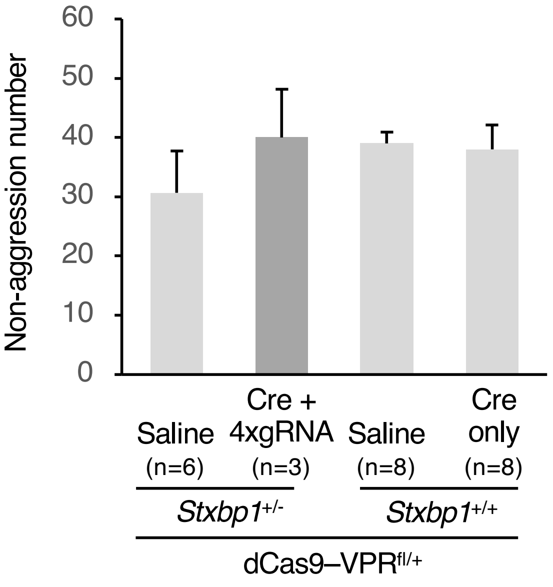


**Supplementary Figure S3. CRISPR-ON treatment did not affect non-aggressive behaviors**. Quantification of non-aggressive behaviors in the resident-intruder test. No significant differences in the frequency of non-aggressive behaviors were detected among the groups.

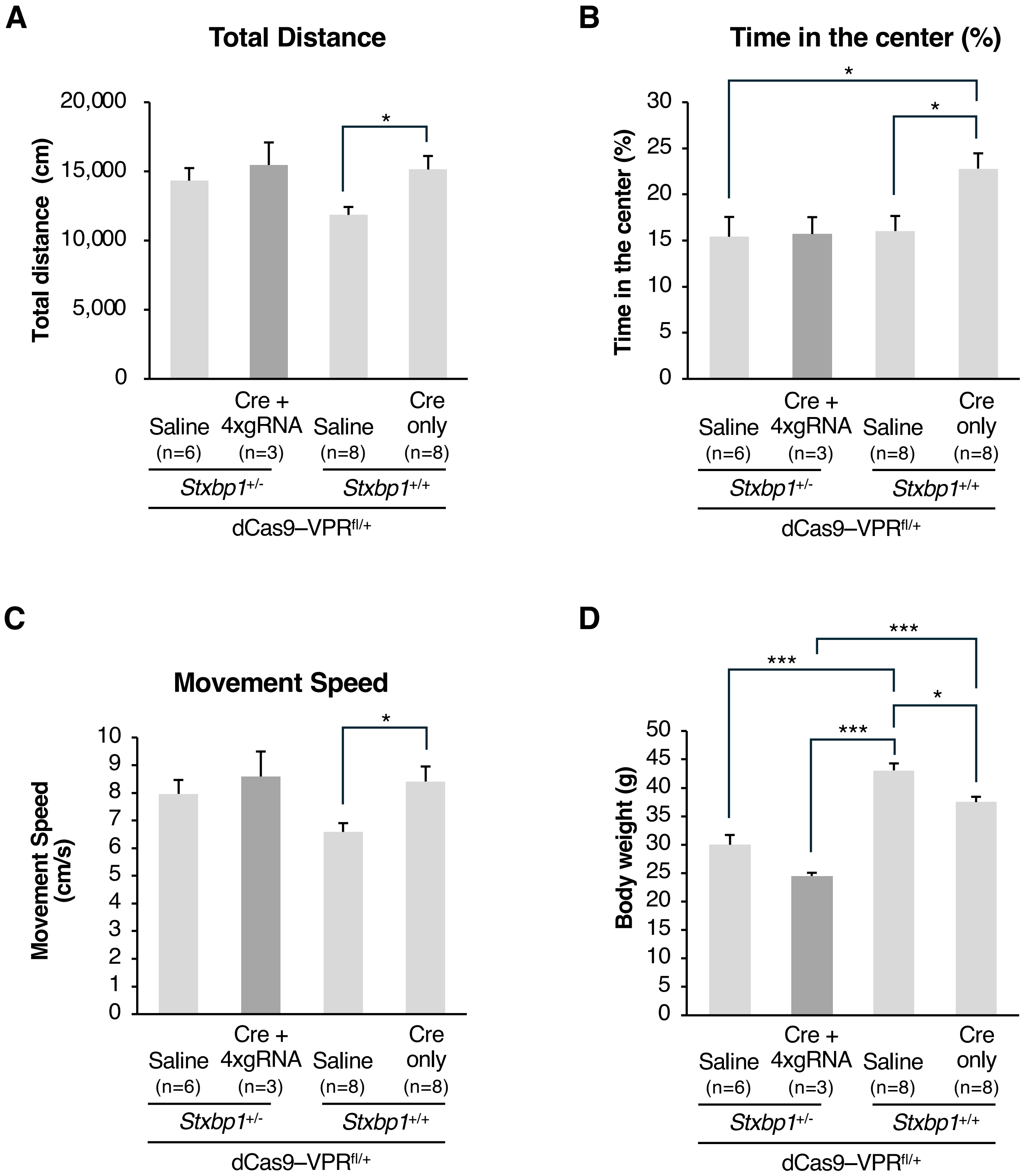


**Supplementary Figure S4. CRISPR-ON treatment does not affect locomotor activity or rescue reduced body weight in *Stxbp1*$^{+/–}$ mice.** (**A–C**) Open field test (10 weeks of age; 30-min sessions) measuring total distance traveled (A), time spent in the center zone (B), and locomotor velocity (C). CRISPR-ON–treated mice did not differ significantly from either untreated *Stxbp1*$^{+/–}$ mice or control mice (*Stxbp1*$^{+/+}$/dCas9-VPR$^{fl/+}$ administered saline) in any of these measures. *Stxbp1*$^{+/–}$ mice showed a trend toward increased distance traveled and faster locomotion compared with control mice. In contrast, mice expressing dCas9-VPR alone (Cre only) showed significant increases in distance traveled, center-zone occupancy, and locomotor velocity compared with control mice. (**D**) Body weight measured at 16 weeks of age. Both CRISPR-ON–treated and untreated *Stxbp1*$^{+/–}$ mice showed significantly lower body weight than control mice, indicating that CRISPR-ON treatment did not rescue the reduced body weight phenotype. Statistical comparisons were performed using one-way ANOVA followed by Tukey's post-hoc test (*$p < 0.05$, ***$p < 0.001$).
